## Supplementary material for "β-arrestin-dependent and -independent endosomal G protein activation by the vasopressin type 2 receptor": Figure 1-figure supplement 1

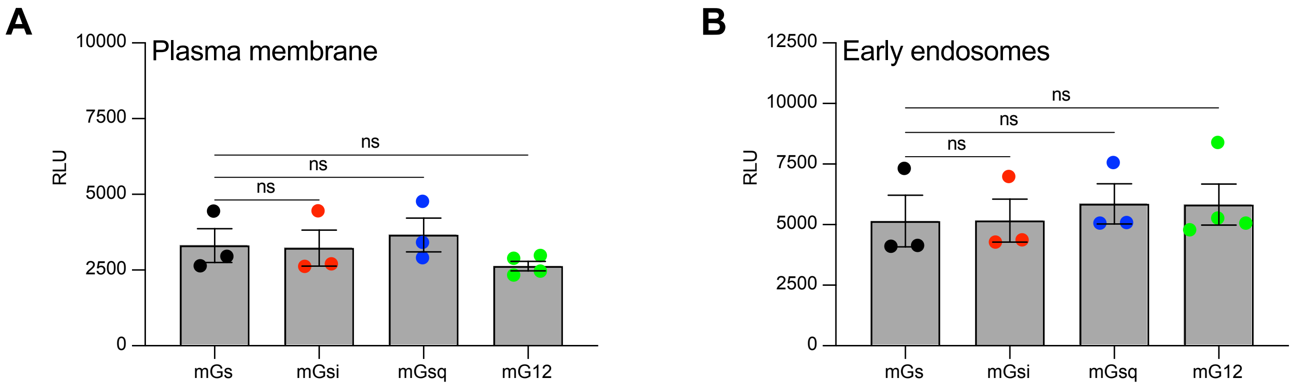


**Equivalent expression of mG constructs for the V$_2$R kinetics**
**(A)** Relative luminescence emitted by each of the four Rluc-fused mG constructs expressed by the cells used to monitor the kinetics of the recruitment of mG proteins at the plasma membrane upon V$_2$R stimulation. **(B)** Relative luminescence emitted by each of the four Rluc-fused mG constructs expressed by the cells used to monitor the kinetics of the recruitment of mG proteins at the early endosomes upon V$_2$R stimulation. $n$ = 3 independent experiments for mGs, mGsi, and mGsq, and $n$ = 4 for mG12. No statistical difference (ns) was detected between the expression of mGs and the other mG proteins using one-way ANOVA and Dunnett's post hoc test for multiple comparisons.
