## Supplementary material for "β-arrestin-dependent and -independent endosomal G protein activation by the vasopressin type 2 receptor": Figure 3-figure supplement 2

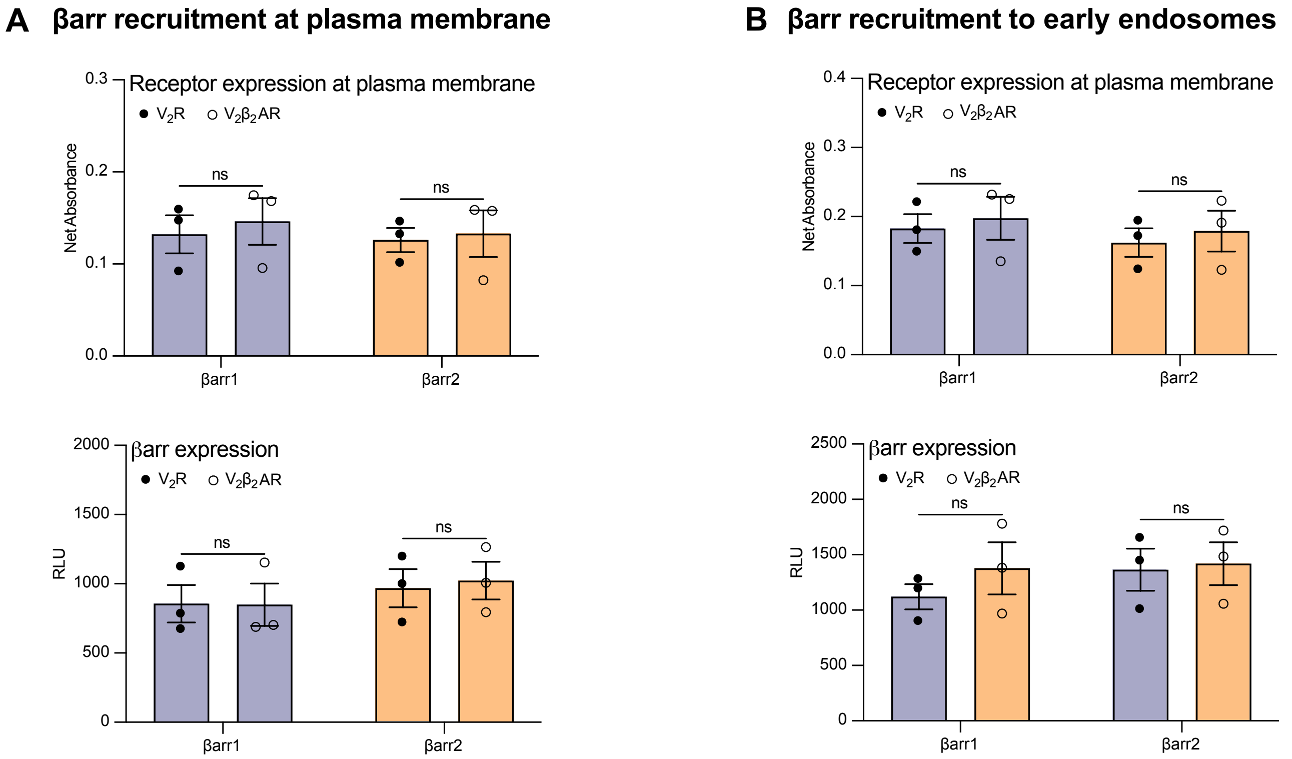


**Relative expression of V2R and V2β2AR at the plasma membrane and of βarrs**

**(A)** Relative V2R and V2β2AR expression at the plasma membrane (upper panel) and of βarr1 and βarr2 (bottom panel) in the βarr recruitment at the plasma membrane experiments. **(B)** Relative V2R and V2β2AR expression at the plasma membrane (upper panel) and of βarr1 and βarr2 (bottom panel) in the βarr recruitment at the early endosomes experiments. *n* = 3 biological replicates for all conditions. No statistical difference (ns) was detected between V2R and V2β2AR using two-way ANOVA and Sidak's post hoc test for multiple comparisons.
