## Supplementary material for "β-arrestin-dependent and -independent endosomal G protein activation by the vasopressin type 2 receptor": Figure 4-figure supplement 1

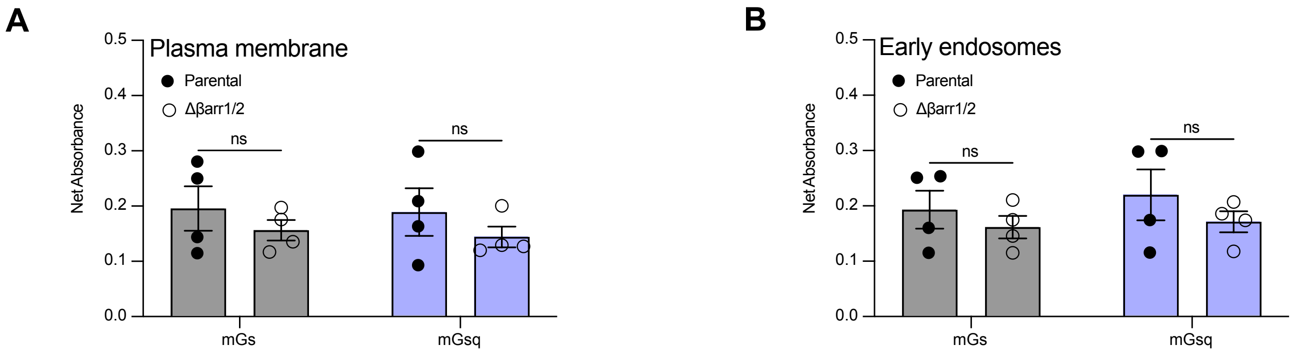


**Relative expression of $V_2R$ at the plasma membrane in parental and Δβarr1/2 cells**

**(A)** Relative expression of $V_2R$ in parental versus Δβarr1/2 cells determined by ELISA in the AVP dose-response curves of mGs and mGsq recruitment to the plasma membrane. **(B)** Relative expression of $V_2R$ in parental versus Δβarr1/2 cells determined by ELISA in the AVP dose-response curves of mGs and mGsq recruitment to the early endosomes. $n$ = 4 biological replicates for all experiments. No statistical differences (ns) were detected between parental and Δβarr1/2 cells as assessed by two-way ANOVA and Sidak's post hoc test for multiple comparisons.
