## Supplementary material for "β-arrestin-dependent and -independent endosomal G protein activation by the vasopressin type 2 receptor": Figure 4-figure supplement 2

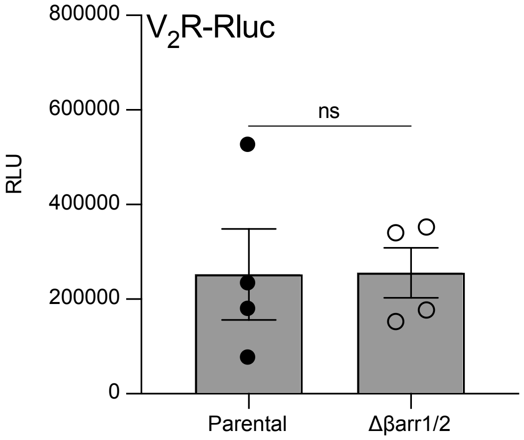


**Relative expression of $V_2$R-Rluc in parental and Δβarr1/2 cells**

Relative expression of $V_2$R-Rluc in parental versus Δβarr1/2 cells determined by monitoring the relative luminescence units (RLU) emitted by Rluc in the kinetics of $V_2$R internalization. $n$ = 4 biological replicates. No statistical differences (ns) were detected between parental and Δβarr1/2 cells as assessed by a paired t test.
