## Supplementary material for "β-arrestin-dependent and -independent endosomal G protein activation by the vasopressin type 2 receptor": Figure 4-figure supplement 3

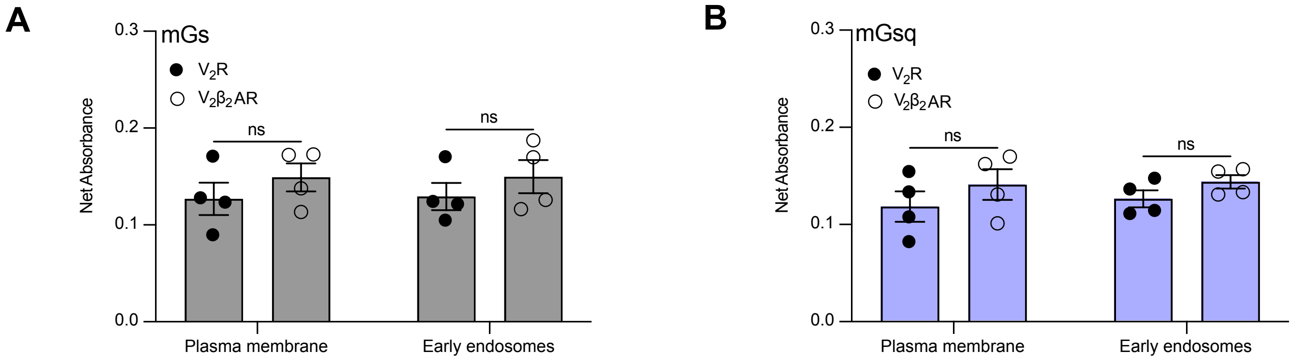


**Relative expression of $V_2R$ and $V_2\beta_2AR$ at the plasma membrane.**

**(A)** Expression of the receptors at the plasma membrane in the cells used to determine the transduction coefficients of the recruitment of mGs at the plasma membrane and early endosomes determined by ELISA. **(B)** Expression of the receptors at the plasma membrane in the cells used to determine the transduction coefficients of the recruitment of mGsq at the plasma membrane and early endosomes determined by ELISA. *n* = 4 biological replicates for all conditions. No statistical differences (ns) were detected between the expressions of $V_2R$ and $V_2\beta_2AR$ as assessed by two-way ANOVA and Sidak's post hoc test for multiple comparisons.
