## Supplementary material for "β-arrestin-dependent and -independent endosomal G protein activation by the vasopressin type 2 receptor": Figure 4-figure supplement 4

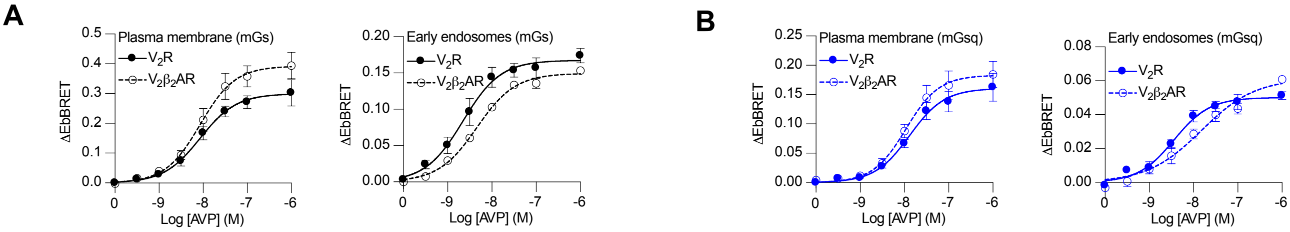


**AVP dose-response curves of the recruitment of mGs and mGsq to the plasma membrane and early endosomes by the V$_2$R and V$_2\beta_2$AR**

**(A)** Dose-dependent recruitment of mGs at plasma membrane (left panel) or early endosomes (right panel) in cells expressing V$_2$R or V$_2\beta_2$AR upon 10 minutes (plasma membrane) or 45 minutes (early endosomes) of AVP treatment. **(B)** Dose-dependent recruitment of mGsq at plasma membrane (left panel) or early endosomes (right panel) in cells expressing V$_2$R or V$_2\beta_2$AR upon 10 minutes (plasma membrane) or 45 minutes (early endosomes) of AVP treatment. *n* = 4 biological replicates for each condition.
